# Droplet microfluidics forward for tracing target cells at single-cell transcriptome resolution

**DOI:** 10.1101/2022.09.13.507865

**Authors:** Yang Liu, Shiyu Wang, Menghua Lyu, Run Xie, Weijin Guo, Ying He, Xuyang Shi, Yang Wang, Jingyu Qi, Qianqian Zhu, Hui Zhang, Tao Luo, Huaying Chen, Yonggang Zhu, Xuan Dong, Zida Li, Ying Gu, Feng Mu, Longqi Liu, Xun Xu, Ya Liu

## Abstract

The rapid promotion of single-cell omics in various fields has begun to help solve many problems encountered in research including precision medicine, prenatal diagnosis, and embryo development. Meanwhile, single-cell techniques are also constantly updated with increasing demand. For some specific target cells, the workflow from droplet screening to single-cell sequencing is a preferred option, which also should reduce the impact of operation steps such as demulsification and cell recovery. We developed an all-in-droplet method integrating cell encapsulation, target sorting, droplet picoinjection, and single-cell transcriptome profiling on chips to achieve a labor-saving monitor of TCR-T cells. As a proof of concept, in this research, TCR-T cells were encapsulated, sorted, and performed single-cell transcriptome sequencing (scRNA-seq) by injecting reagents into droplet. It avoided the tedious operation of droplet breakage and re-encapsulation between droplet sorting and scRNA-seq. Moreover, convenient device operation will accelerate the progress of chip marketization. The strategy achieved an excellent recovery performance of single cell transcriptome with a median gene number over 4000 and a cross-contamination rate of 8.2 ± 2%. Furthermore, this strategy allows us to develop a device with high integrability to monitor infused TCR-T cells, which will promote the development of adoptive T cell immunotherapy and their clinical application.

## Introduction

Since the implementation of the Human Genome Project in the last century, the popularization of DNA and RNA sequencing technologies showed tremendous impact on the development of the entire life sciences and medical fields. With the deepening of research, we find substantial differences between thousands of cells in a tissue, and even phenotypically identical cells also have heterogeneity and differences in gene expression levels^1, 2^. Therefore, the interpretation of single-cell genetic information provides stronger technical support for cancer treatment, embryonic development, prenatal diagnosis, and so on^3, 4^. Because of this, a revolution in single-cell sequencing may rebuild the cognition of cellular life. The rapid evolution of single-cell omics is inseparable from the fast upgrade of application tools. In 2015, Drop-seq was proposed to set up a bridge between single-cell profiling and high-throughput technology^5^, and also laid the foundation for the development of similar tools at a later stage. The introduction of droplet microfluidics also directly stimulates the throughput of single-cell sequencing to increase exponentially, and RNA profiling of millions of cells has been achieved in less than a decade^6^.

Without exception, under the mainstream single-cell tools with microfluidics as the technical background^7, 8^, experimental samples require processes such as single-cell suspension preparation, droplet generation, mRNA capture, demulsification, and so on. However, in some specific application, there is a need to enrich target cells with specific needs from the sample in advance, the subsequent single-cell sequencing process will also have strict requirements on cell input and viability. For instance, engineered T cells equipped with receptors can specifically recognize the antigens on tumor cells^9, 10^, and then initiate cytotoxic effects to suppress tumors. Sorting and unveiling the phenotypes of T cells pre- and post-infusion will be instructive to uncover their cellular biology and microenvironmental factors underpinnings. Alternatively, by integrating plasma cell encapsulation, antibody secretion screening in droplets, and scRNA-seq into a pipeline on chips, the time for identifying antigen-specific antibodies is compressed to 24 hours from weeks ^11, 12^. Commercialized fluorescence-activated cell sorting (FACS) is preferred for enriching cells and allowing samples to adapt microwell ^13^ or droplet-based RNA-sequencing ^14^. To promote efficiency and convenience, droplet-based microfluidics systems have been developed to handle the sorting and single-cell analysis. Recently, dielectrophoresis array ^15^ and differential flow resistance principle ^16^ improve cell utilization rate of micro-well chips, and increase the capability of handle scRNA-seq for abundant cells. However, the absence of sorting function makes these methods not always appropriate for target cell scRNA-seq. As a comparison, droplet-based single-cell sequencing has achieved huge breakthrough in many areas, thanks to its high convenience in operation. Importantly, similar florescence-based optical sorting methods have been available for years and can induce high efficiency. However, a drawback of existing droplet-based strategies is the inevitable step of sorted droplet breakage and re-encapsulating cells for downstream single-cell sequencing, which would affect the cell viability and total number ^11, 12^.

Herein, we integrated sample preparation based on droplets, target sorting, droplet picoinjection, and single-cell transcriptome profiling to generate an efficient platform for examining TCR-T cells at single-cell resolution (**Figure 1A**). As a proof of concept, TCR-T cells in a mixture are labeled with fluorescence, and then cells are co-encapsulated with RNA-beads (DNBelab C4) for transcriptome information capture in droplets. For enrichment of TCR-T cells, targets are sorted into a collector which is compatible with the pico-injector inlet hole based on the activation of fluorescent tags. Negative pressure is transiently applied to the chip, and droplets are driven to the flow channel, where reagents for scRNA-seq are injected into droplets facilitated by an electric field. Afterward, the cells were subjected to lysis inside droplets, transcripts capture, library preparation and RNA-sequencing.

**Figure 1.**
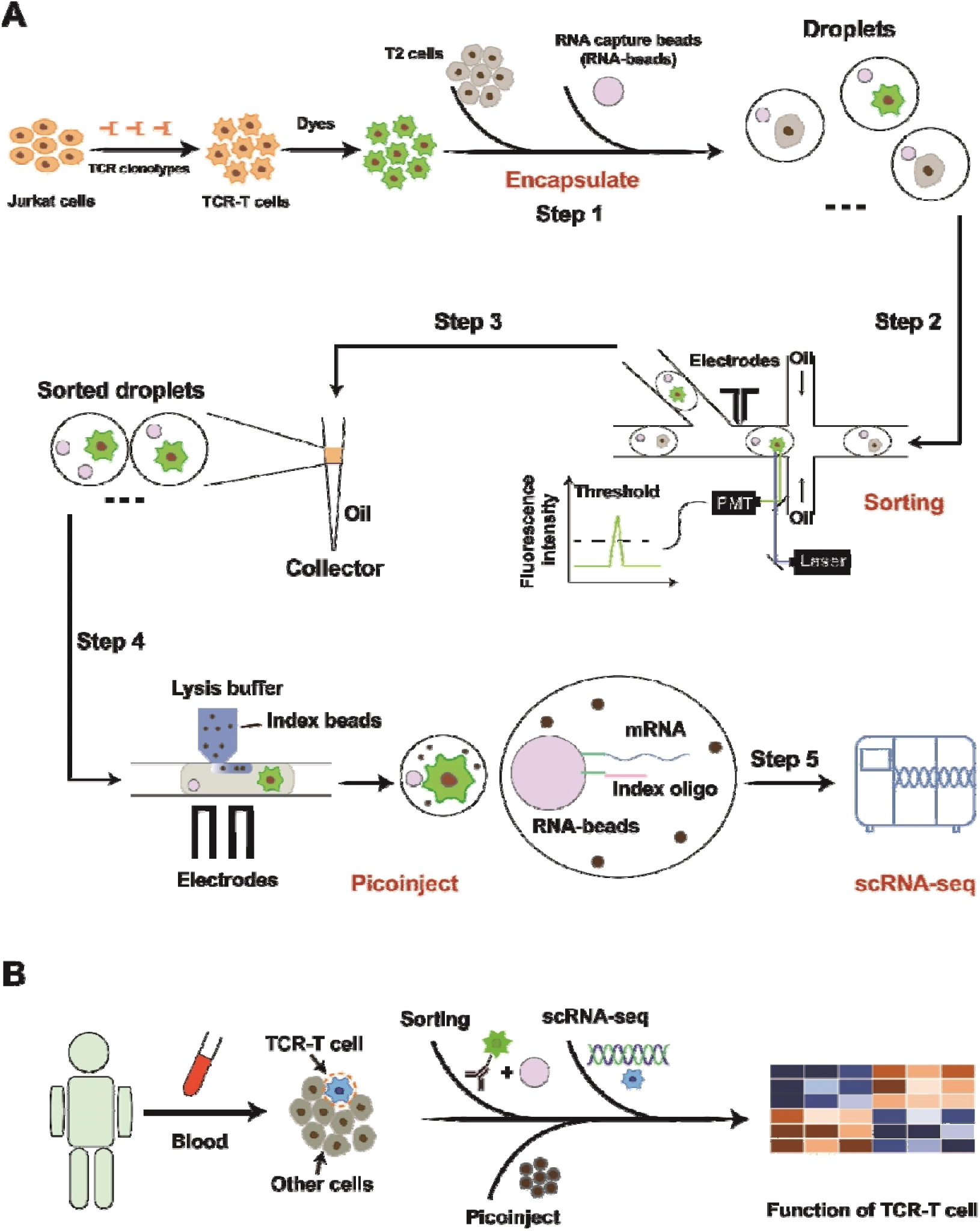
Schematic diagram of experimental flow. A) Step1: Droplet generation: Co-encapsulation of cells and RNA-beads. Step2: Droplet sorting: Fluorescence-activated droplet screening and sorting for enriching TCR-T cells. Step3: droplet collection; Step4: Droplet picoinject: Inject reagent (lysis buffer and index beads) into sorted droplet for scRNA-seq. Step5: Single-cell RNA-sequencing: Capture transcripts, prepare library and perform scRNA-seq. B) The strategy for monitoring infused TCR-T cells in clinic application.

The pipeline achieves a high-throughput cell sorting rate (up to 450 droplets per second), convenient processes without steps of breaking and re-generating droplets, and ideal quality of single-cell transcriptome (median gene number of per cell: ~4000). Then, the cell-cycle phases and other phenotypes of TCR-T cells were presented in a demo, highlighting the potentiality of our strategy for unveiling heterogeneous phenotypes of TCR-T cells. The droplet injection technology involved in this work is actually a crucial step towards the development of the new method for antigen-specific TCR screening. In previous work, we reported the utilization of droplet microfluidics to sort antigen-specific T cells at a single-cell level, which is conducive to the rapid search for TCRs targeting neoantigens ^17^. In that study, we must perform demulsification and cell recovery after target screening before the next step of sequencing work. In the identification step, emulsion-breaking will cause huge obstacles to the identification and backtracking of pairing relationships (TCR-Neoantigen). So, we proposed a novel method of linking droplet injection and single-cell omics to realize it. It also lays the foundation for establishing the recognition mechanism of tumor neoantigens and their specific TCRs from the single-cell transcriptome level. In the future, the strategy can be used for clinical samples to uncover phenotypes relating to favorable prognosis after infusions, and provide clues to design more effective therapies (**Figure 1B**).

## Materials and experiments

### Chip design and fabrication

This study employed AutoCAD to design three different microfluidic chips, including droplet generation chip (Chip A), droplet sorting chip (Chip B), and picoinjection chip (Chip C) (**Figure S1**). The master models were fabricated using SU-8 photolithography on 4-inch silicon wafers and the chips were fabricated by PDMS as previous described ^17^. Given the hydrophobic performance ^18^, Chip A was bonded on a PDMS-fabricated substrate, Chip B and C were bonded to glass slides after being exposed to oxygen plasma (Harrick plasma cleaner, PDC-002). The details of chips design and fabrication are listed in the Supporting Information.

### Cell culture

T2, Jurkat, NIH/3T3 (CRL-1658) and 293T cell lines were purchased from American Type Culture Collection. The TCR-T cell line was generated by transducing a TCR lentivirus vector into Jurkat cell line as previously described ^17^. TCR-T cells and T2 cells were cultured in RPMI 1640 supplemented with 10% (v/v) fetal bovine serum (FBS) and 1% penicillin-streptomycin (P/S) at 37 □ with 5% CO_2_. NIH 3T3 and 293T cell lines were cultured in Dulbecco’s Modified Eagle Medium (DMEM) supplemented with 10% (v/v) FBS and 1% P/S at 37 □ with 5% CO_2_.

### Cell staining and flow cytometry

TCR-T Cells were washed with DPBS once and resuspended in 1 mL DPBS with 1 μL CellTracker Green CMFDA Dye. After incubation at 37 □ for 30 minutes, cells were washed 3 times with 1 mL DPBS. Then cells were subsequently centrifuged at 500 g for 3 minutes, and resuspended in working buffer, comprising RPMI 1640 supplemented with 0.1% Pluronic F-68 (diluted from 10% Pluronic F-68), 1% P/S, 100 mM tris(3-hydroxypropyl) phosphine (THPP) and 12% Ficoll PM400 at final density of 1000/μL. For flow cytometry (Aria II, BD, USA), cells were washed with DPBS twice, centrifuged at 500g for 3 minutes and resuspended in DPBS with 2% FBS.

### Single-cell RNA sequencing

The droplets were incubated for 40 minutes at room temperature, and then be demulsified to recover RNA-beads. The cDNA library for scRNA-seq was generated by DNBelab C4 kit ^19^. Then, all libraries were conducted in preparation for DIPSEQ T1 sequencer (MGI). The detailed method for data analyzing and data mining are shown in the Supporting Information.

### Statistical analysis

The statistics were conducted by GraphPad and R. The difference between two groups were examined by Wilcoxon test, and the *p* values of multiple tests in **Figure Figure 5** was adjusted by false discovery rate (FDR) method.

### Data Availability

The data that support the findings of this study have been deposited into CNGB Sequence Archive (CNSA) ^20^ of China National GeneBank DataBase (CNGBdb) ^21^ with accession number CNP0002544.

## Results and discussion

### Platform operation

To achieve TCR-T cell enrichment and scRNA-seq based on chips, droplet generation chip (Chip A, **Figure S1A, B**), droplet sorting chip (Chip B) and picoinjection chip (Chip C, **Figure S1C, D**) were designed and manufactured in this research. Chip A has two inlets for samples and one inlet for oil as well as an outlet for droplets (**Figure S1A**). The outlet hole was connected to a 2 mL collection tube and the negative pressure driver of 30 mL syringe was assembled into a 3D-printed base for supplying force in droplet generation. Then, 50 μL of bead/cell solution and 400 μL of oil were added into the individual holes (**Figure S1A**). The negative pressure of ~13kpa as the syringe is pulled up can induce a transient and stable droplet generation, and meanwhile the overabundant residual in sample and bead/cell sedimentation in the hole could be effectively avoided. The time for generating millions of droplets was less than 10 minutes, and no more than 6% pressure decline under the supervision of sensors (4AM01, LabSmith, USA) could be contributed (**Figure S2A**).

The Chip B was adapted from previous published design ^17^,the droplets and oil input were driven by two independent syringe pumps, and the flow rate was set to 80 μL and 800 μL per hour, respectively. When a target droplet was identified, a 20 kHz of 1000 peak-to-peak voltage (~2 ms duration) would be applied to electrodes for cell sorting.

In Chip C, one outlet and three inlets were punched for inputting droplets, oil, and injected reagent, respectively. The entire infusion procedure took approximately 20 minutes by pulling the syringe from 17 mL to 20 mL scale, and the negative pressure decay rates were controlled to be with 5% (**Figure S2B**). In this system, the injection leakage is an unavoidable phenomenon. The pressure differential between the injector and the oil channel is approximately determined by Laplace pressure ^22, 23^. However, the picoinjector used herein was driven by negative pressure, suggesting that the pressure in injector and oil channel were approximately equal to atmosphere pressure and no differential value was generated. Therefore, this situation eventually led to the injection leakages. Instead, connect injection to a syringe pump may be a feasible manner which could balance the Laplace pressure in turn.

### Cell encapsulation, sorting, and injection

TCR-T was firstly labeled with CellTracker CMFDA and mixed with T2 cells to mimic cell-type proportions in blood. Then, the CMFDA stained cells would generate a signal for subsequent droplet sorting. Since the diameter of droplet for efficiently sorting is required to be 40-60 μm. To overcome the influence of Poisson distribution as much as possible, the droplet size generated by Chip A was controlled to be ~60 μm. According to the statistics of 7737 droplets, the average diameter of droplets is 56.87 ± 1.95 μm (**Figure S3A**). In order to verify the stability of the droplets, we put the generated droplets into the incubator for 4 hours and 6 hours, and then counted the diameter distribution of the droplets. **Figure S3B, C** showed that the droplets did not change significantly in 4-6 hours. In this research, subsequent experimental operations could be carried out without incubation after droplet generation, but incubation may be required for 4-6 hours after droplet generation in the future work. To minimize the possibility of more than one cell in a droplet, cell concentration was adjusted according to Poisson distribution with λ = 0.05. In **Figure S3D**, the experimental cell distribution in droplets was approximately determined by Poison distribution. Therefore, when the cell density was adjusted to 1000/μL, the rate of droplets with one cell was 4.91 ± 0.16%, and that with more than one cell was 0.13 ± 0.02%, suggesting that multiplet rate was less than 5%. For single-cell sequencing data, 5% is an experience value to avoid the influence of multiplet on cell omics profile ^24^ In terms of improving cell utilization, RNA-beads were over-loaded (25000/μL). Our study showed that the percentage of droplets without RNA-bead(s) was about 25% at the bead density of 25000/μL. As shown by **Figure S3D**, the proportion of droplets containing RNA-bead(s) was 74.56 ± 0.53%, and 3.70 ± 0.04% of droplets simultaneously contained RNA-bead(s) and cell(s), suggesting that 73.42 ± 1.59% of droplets encapsulating cells also had beads within (the gray column). Given that only RNA captured by beads could be performed in scRNA-seq, this operation helped us to decrease the loss of cells.

Droplets containing target cells were enriched by the fluorescence-activated droplet sorting (FADS), and in **Figure 2A**, fluorescence signal (green) from the CellTracker CMFDA was then accepted and transduced into voltage signal by photomultiplier tube (PMT). Additionally, the voltage was simultaneously delivered to comparator and single chip microcomputer (SCM). If a voltage exceeded the threshold of the comparator, a high voltage (~1Vpp) operating on the full cycle of the PMT signal would be transmitted from the comparator to SCM for triggering PMT signal recording. After the comparator signal ended, the SCM calculated the maximum value of the PMT signal and compared it with the threshold. Thus, a PMT signal above threshold sent a voltage to sort the droplet by dielectrophoresis.On the contrary, cell-free droplets or non-fluorescent-cells did not excite signal peaks (**Figure 2B**). The approach of droplet sorting finally leaded to a percentage of positive cells over 95% re-assessed by flow cytometry (**Figure S4**), whereas 35% corresponds to the percentage before sorting (**Figure 2C**).

**Figure 2.**
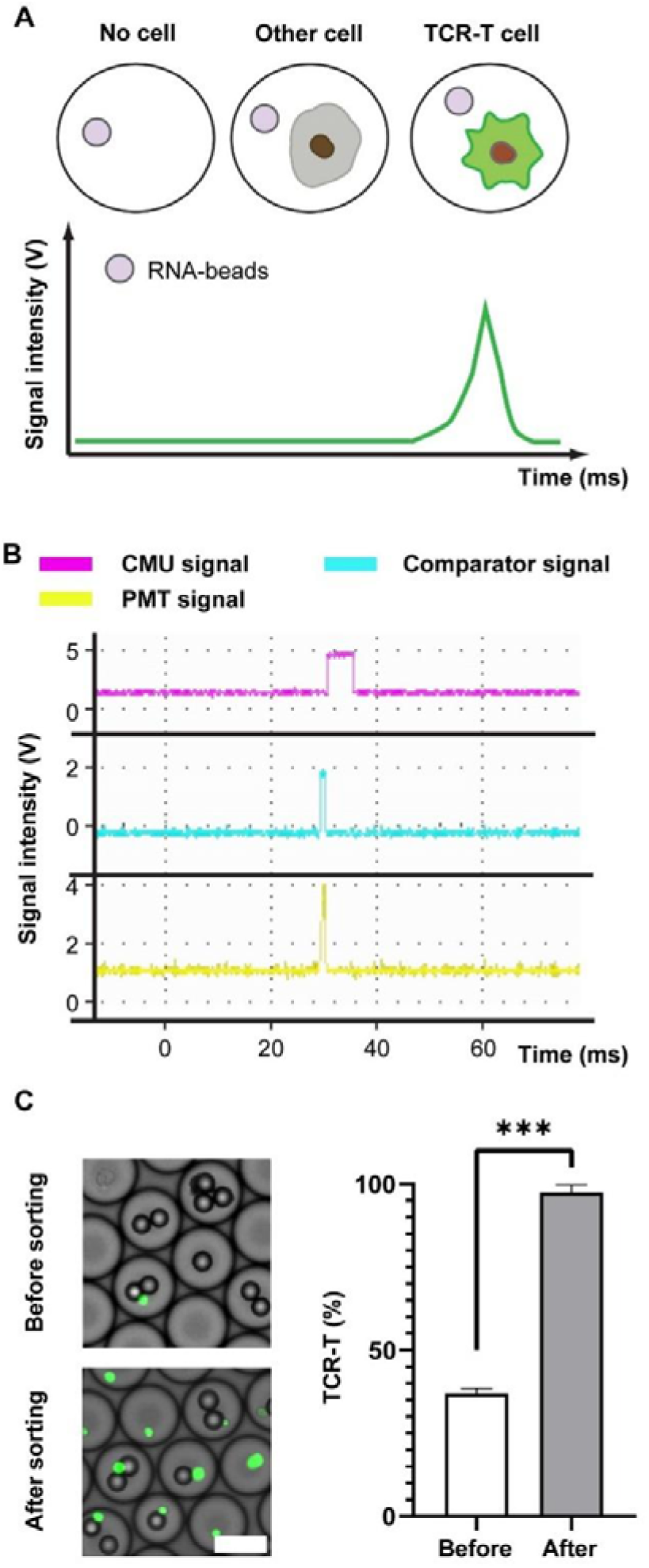
Droplets sorting and signal analysis. A) The signaling patterns of droplets with cell-free, off-target -cell or TCR-T cell. B) A signal recording of droplet with TCR-T cell, and feedback signals from comparator and CMU against a positive droplet. C) Images of droplets before and after sorting (left), and the percentage of CMFDA+ cells detected by flow cytometric analysis. The TCR-T cells are highlighted by green points. Scale bar: 50μm. Wilcoxon test is used, and *\*\*\*p* < 0.001.

We connected the outlet catheter to a 200 μL pipette tip, and added 20 μL of empty droplets to prevent further cell loss. Owing to the limited carrying capacity of tip, the tip was inserted to a device of which pressure be operated by a 1 mL syringe for controlling the liquid level, and the detailed procedure was presented in **Figure S5.** However, there is a risk that the transfer process may influence the uniformity of the droplets, and we also conducted stability statistics of droplets during the transfer from Chip B to Chip C. As we can see from **Figure S6A, B**, droplet transfer operations have no impact on droplet stability. After sample collection, the pipette tip was then inserted into the inlet on Chip C (**Figure 3A**), and index beads were re-suspended in the lysis buffer at a density of 70,000/μL as the injection phase to ensure that each droplet contained index beads after injection. In **Figure 3B**, droplets and inject solutions were dyed with red and green pigments to facilitate observation, respectively. Following completion of color-dye injection, the orange color of droplet reflected a given action. Two distributions of droplets were presented in **Figure 3C**, in which the uniform values of small (green, 51.22 ± 3.51 μm) and large ones (orange, 79.31 ± 6.23 μm) indicated the stability of injector. After incubation at room temperature for 40 minutes in the collection tube, cells in droplets were lysed, index was cut by reagent, and released mRNA and index were captured by RNA-beads (**Figure 3D**). In order to further explore the stability of droplets, we made statistics on the diameter of droplets before and after incubation (**Figure S6C, D**). Through statistical calculation, it was found that the diameter distribution of droplets did not change significantly after 40 minutes of incubation, which confirmed that the incubation process had no effect on droplet stability. Finally, with droplets breakage, RNA-beads were recovered for library preparation as previously described ^17^. **Figure S7A** showed the principle of nucleic acid capture by RNA-beads, and **Figure S7B** reflected that the distribution of cDNA and oligo after separation and purification, which is consistent with the results of conventional library construction. This indicated that the library can be used for subsequent single-cell sequencing and data analysis.

**Figure 3.**
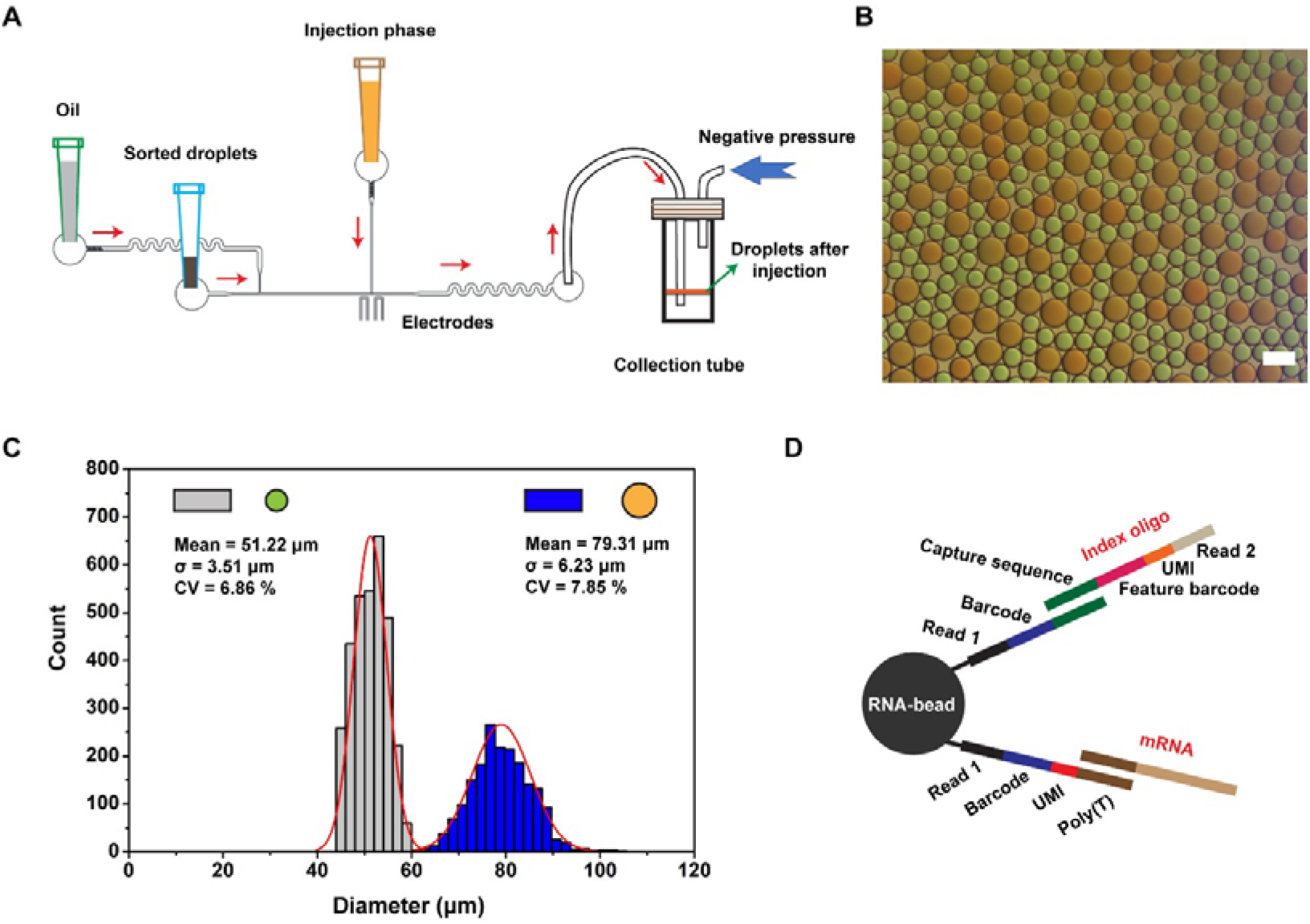
Droplet picoinjection and oligo design for RNA-beads. A) The schematic diagram of droplet picoinjection. Negative pressure drives oil, droplets, and injection reagent to flow through channels. Oil control the distance of two adjacent droplets, and additional reagents are injected into droplets facilitated by an electric field. B) The photograph of droplets after injection. To facilitate observation, droplets and injection were dyed with red and green, respectively, and then red droplets were orange, and leakage droplets of injection were green after injection. Scale bar is 100μm. C) Diameter distribution of droplets (N = 3203, grey bar; N = 1870, blue bar). D) RNA and index are captured by RNA-beads with different capture handles. Sequence structure for capturing index: bead, read1, barcode and capture sequence; for capturing RNA: bead, read1, barcode, UMI and poly(T).

### Data analysis for single-cell sequencing

3000 and 5000 TCR-T cells were sequentially sorted for picoinjection and scRNA-seq library preparation. After accomplishing the above-mentioned steps, the raw data were aligned and annotated by the PISA pipeline. After normalizing sequencing depth to that of 10X V3 demo, the gene number of 3000- and 5000-cell samples were 3484 ± 300 and 4046 ± 154, respectively, and the two groups (3000 and 5000-cell samples) did not have obvious differences in UMI and mitochondrial genes rate (**Figure 4A**). Thereby, the factor of cell number did not affect the genes detection in this method, and gene number detected here was better than the majority of common scRNA-seq methods ^25, 26^ (**Figure 4B**). However, the recovery rate of 3000-cell was less than that of 5000-cell samples (**Figure 4A**). To further explore the problem of cell recovery rate, we have provided a set of experiments (1000-cell samples). In **Figure S8** and **Figure 4A**, we found that the trend was not consistent when the cell input was reduced to 1000. This is because when the number of samples is reduced, the RNA-beads input would also be decreased. In subsequent operations, experiments that rely on magnetic bead recovery may cause fluctuations in cell recovery rates due to some experimental errors. To overcome these limitations in the future, performing reverse transcription-PCR^27,28^ in droplets and/or replacement of RNA-beads with dissolvable material ^29, 30^, and then recovering RNA or cDNA solution may further increase the recovery rate.

**Figure 4.**
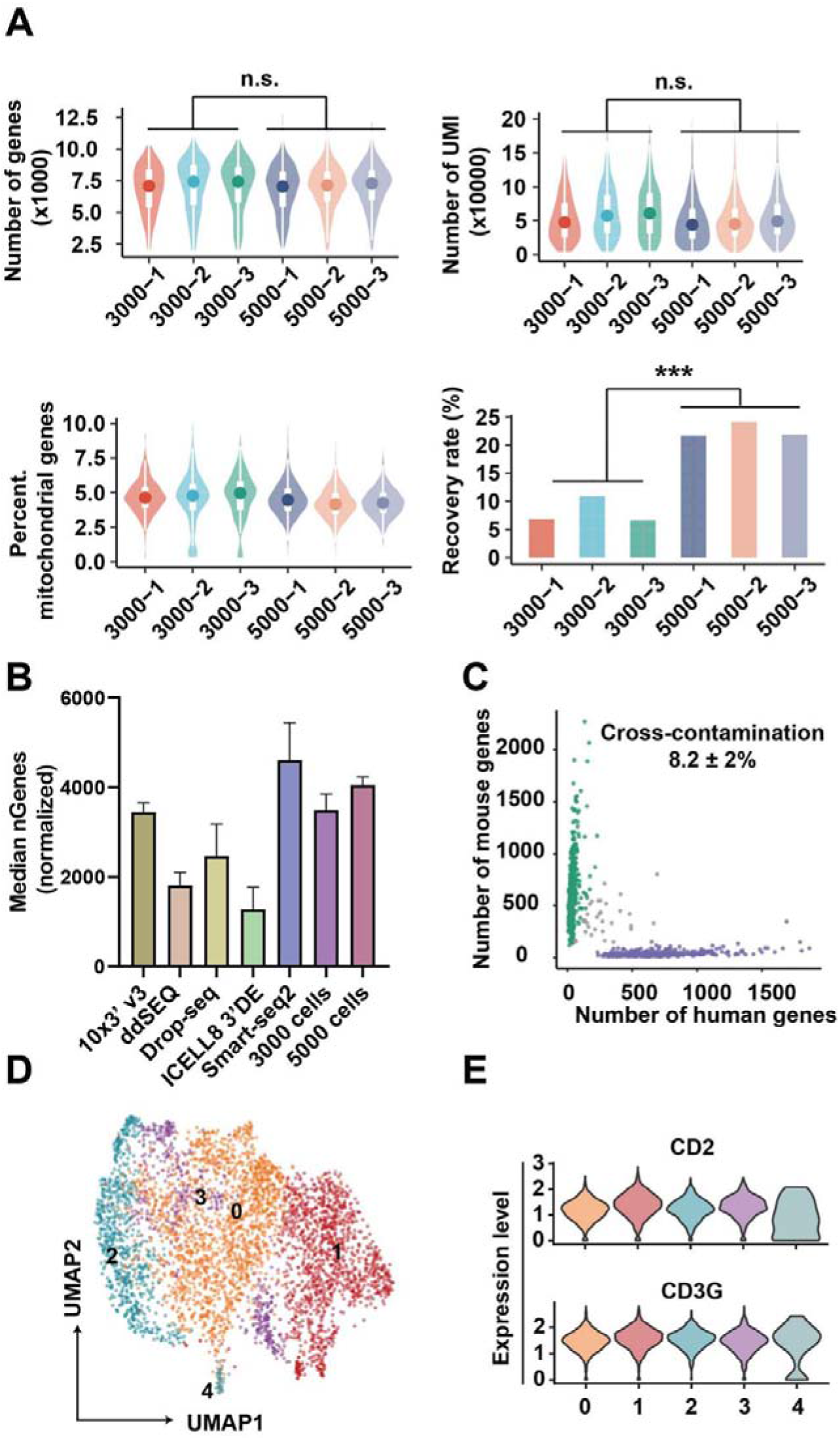
Quality of scRNA-seq data. A) From the upper to lower, the number of genes per cell (1st), UMI per cell (2nd), and the percentage of mitochondrial genes per cell (3rd), and cell recovery rate (4th). B) The median of genes per cell generated by other methods and our method. The gene numbers of 10X, ddSEQ, Drop-seq, ICell8 and our method were adjusted by read number to normalize sequencing depth. C) The number of genes of human or mouse in each cell. D) UMAP of cell clusters when SNN graph was construed with a resolution = 1. E) Features markers of TCR-T cells.

**Figure 5.**
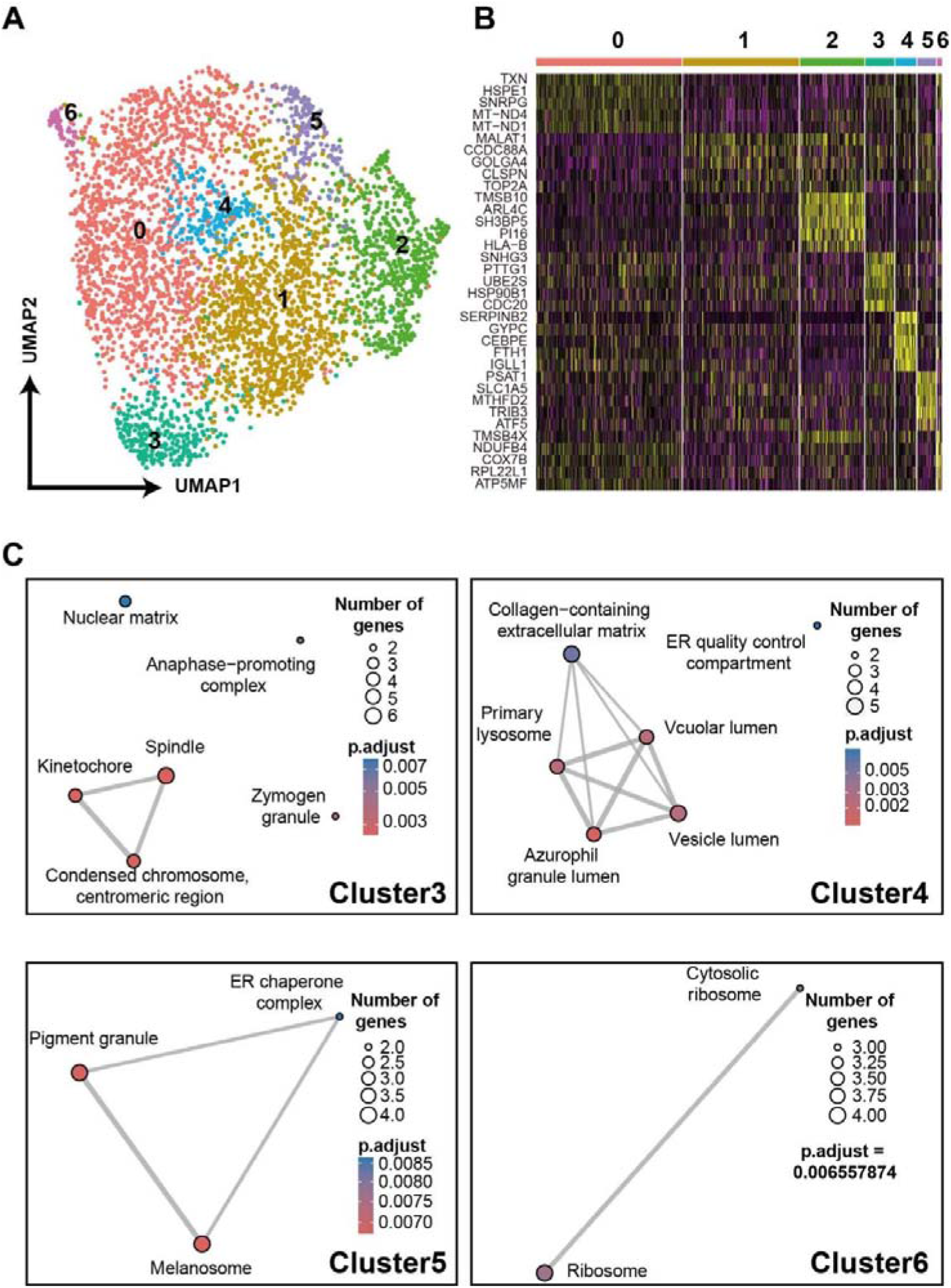
A demo for unveiling TCR-T phenotypes with scRNA-seq data. A) UMAP of TCR-T cells after cell-cycle regression. B) Heatmap of top5 DEGs of each subset. C) GO analyses based on DEGs of each subset.

To estimate the cross-contamination, 293T (human origin) and NIH 3T3 (mouse origin) cells were equally mixed together for scRNA-seq by our method, and the cross-contamination rate in **Figure 4C** was 8.2 ± 2%. According to previous reports, the cross-contamination detected by scRNA-seq data was biased from the theoretical value predicted by the Poisson distribution ^31^. As shown by Yang et al. ^32^, ambient RNA released in the cell suspension is a notable factor to increase the cross-contamination. Likely, undergoing apoptosis and tressed cells may release these RNA molecules to cell suspension, which are further incorporated in droplets. Beyond ambient RNA, other experimental factors, such as barcode swapping may also cause cross-contamination ^33^. Reducing cell concentration should be efficient to repress multiplet, and some algorithms can facilitate the detection of varied types of pollution ^32, 33^. In addition, cells from all samples were distributed evenly on a two-dimension UMAP plot, and the result in **Figure S9** reflected that batch effects were adjusted sufficiently. Cells were grouped into five clusters with a resolution = 1 of SNN (**Figure 4D**), and strong CD2 and CD3G expressions in all clusters (**Figure 4E**) demonstrated that features of cellular transcriptome profile can be preserved by this method.

TCR-T cells *in vivo* can present heterogeneous phenotypes due to their diverse phenotypes pre-infusion ^34^ and the various effects of microenvironment after infusion ^35^. Next, we further examined gene expressions of TCR-T cells indicated by scRNA-seq data proving that it can uncover the heterogeneity of TCR-T cells and the underlying mechanism. By DEG analysis, the top10 feature genes of each cluster were presented in a heatmap (**Figure S10A**). The *MCM4, PCNA, RRM2* were highly expressed by cluster0, *CCNB1* and *CDC20* were over-expressed by cluster1, and *CDKN2D* was highly expressed by cluster3. All of these genes are related with cell cycle. According to the analysis of cell-cycle feature genes (**Figure S10B**), the result reflected cells were at different phases (**Figure S10C**). Then, data of cells were scaled again after a regression of cell cycle genes (**Figure S10D**). For **Figure 5A**, cell clusters were skewed and presented novel features (**Figure 5B**). With annotation of DEGs by GO analysis, cluster3 expressed spindle and kinetochore related genes, implying that cells in this cluster were ongoing cell differentiation, and vesicle lumen and vacuolar lumen related genes were highly expressed in Cluster 4. As reviewed by Gutiérrez-Vázquez et al. ^36^, these lumen systems tightly regulated the exosome and other cellular transportation systems, which in turn affected the cell-cell interactions and cellular functions. The DEGs in cluster5 were involved in melanosome, and those in cluster6 were participated in ribosome functions. Results of these analyses on DEGs unveiled the heterogeneously cellular functions of TCR-T cells, and suggested that our method was sufficient to trace TCR-T cell phenotypes. Alternatively, complicated environments and cell-cell interactions in clinical practices will induce a more diverse phenotypes of TCR-T cells *in vivo* ^37^, and our method could transfer more message to uncover the skewed functions of infused cells, which in turn provide clues to develop the adoptive T cell therapies.

In terms of current studies on TCR-T/CAR-T *in vivo*, the cell expansion, exhaustion, and trafficking are primary parameters used for evaluating the therapy responses, which can be obtained by flow cytometry and cell imaging analyses ^37^. Of course, these parameters were also presented by scRNA-seq in our method. Importantly, other effects such as cell cytotoxicity and cytokines secretion can also be exhibited by single cell omics, which would provide more reliable information to generate accurate model for estimating TCR-T/CAR-T treatment effect. At the beginning of the project, we also considered integrating the different functional units onto one chip. However, the purpose of our platform is not just for one single application scenario, and we expect that several chips with different roles may be freely combined like building blocks to achieve the best effect. The next application point is to rapidly identify the pairing relationship between TCRT and APC cells (**Figure S11**). We labeled various of TCR-T and APC cells with different barcodes and then encapsulated these cells in droplets along with RNA-beads. After incubation for a certain time, the APC cell in the droplet would activate corresponding TCR-T cells by presenting specific antigens, which resulting in changes in gene expression of relevant activation pathways. Then, the corresponding barcodes of TCR-T and APC cells were determined by molecular information recovery, library generation and data analysis. Ultimately, we hope to speed up the identification of tumor neoantigens. We also expect that our platform combines the diverse omics approaches (proteomics, epigenetics) to allow a considerable potential for enhancing the development of cell therapy.

### Conclusion

In this study, we developed a strategy with droplets generation, droplet sorting and picoinjection to integrate cell sorting and single-cell omics. The strategy enriched target cells in a high-throughput manner, and performed single-cell omics through injecting reagents into droplets instead of tedious manipulation. This strategy simplifies the pipeline to perform single cell sequencing for enriched cells, and allows a further integration of chip to promote cell biology studies and clinical trials. In the future, we can use this strategy to verify the function of TCR-T cells by taking blood from patients, and achieve monitoring the infusion of TCR-T cells. This step will promote the development and clinical application of adoptive T cell immunotherapy.

## Acknowledgements

We acknowledge financial supports from National Key Research and Development Program of China (2021YFF1200500), Science, Technology and Innovation Commission of Shenzhen Municipality (JSGG20180508152912700), Guangdong Basic and Applied Basic Research Foundation(2021A1515110459), China Postdoctoral Science Foundation (2021M692212), and Shantou University Scientific Research Foundation for Talents (NTF20034).

## Supplementary data

Supplementary data to this article can be found online.

