## Supporting Information for "Droplet microfluidics forward for tracing target cells at single-cell transcriptome resolution"

**Additional Experimental Section**

**Details of chips design and fabrication**

In brief, PDMS base and curing agent were mixed at weight ratio of 10:1, poured onto the mold, and incubated at 80 °C for 1 hour to facilitate the solidification. PDMS chips was peeled off from the model and cut along the chips’ outline. In Chip A, one oil inlet hole and two cell/beads inlet holes were made with hole punchers with 5 mm and 10 mm diameter. In Chip B and Chip C, all inlet and outlet positions were punched into 1 mm diameter holes. And then these microfluidic chips were fabricated by commonly used materials of PDMS and glass, and noted that surface of PDMS is more hydrophobic than glass. Since Chip A is used to generate water-in-oil droplets, the substrate requires sufficient hydrophobicity to stabilize droplets and prevent uncontrolled coalescence. And Chip A was heated for at least 24 h at 60 ℃ to meet hydrophobic performance. Therefore, PDMS layer would be a more appropriate selection for the substrate of the Chip A. As for the Chip B and Chip C, the electrodes were fabricated by infusing melted solder wires into the pre-designed channels on a 90 ℃ hot plate, and then conducted further curation at room temperature. For Chip C, the electrodes were laid at the opposite side of injection across channels for droplet flowing.

**Microscope imaging**

Bright-field images were captured by the Olympus microscope (IX71, Japan) with a high-speed camera (DP26, Olympus, Japan), the pictures of fluorescence-labelled TCR-T cells were acquired using the inverted fluorescence microscope (IX73, Olympus, Japan). Droplets were dropped on a glass slide, and imaged with a 10× objective lens of an inverted microscope. Then, these pictures were automatically quantified by ImageJ software for acquiring statistical data.

**Data analyzing**

The FASTQ data generated by sequencer were processed by PISA pipeline (https://github.com/single-cell-BGI/scRNA-pipe), and a standard format of gene expression matrix was generated with reference to GRCh38 human genome for downstream analysis. The gene expression matrix was processed as input for Seurat software (https://github.com/satijalab/seurat). Genes expressed in < 3 cells and cells with < 200 genes were filtered out. To minimize the batch effect among samples, all samples’ analyses were treated by the reciprocal PCA pipeline (parameter: dims = 20) in Seurat to merge into an integrated dataset. The first 30 PCs were summarized to construct a SNN network, and distance between cells and clusters were calculated by Louvain algorithm. Then, dimensional reduction and illustration of cells in 2-dimension space were performed with UMAP.

**Differential expressed genes (DEGs), gene ontology (GO) enrichment and cell cycle analyses**

Identification of DEGs among clusters by “FindAllMarkers” function in Seurat revealed the heterogeneity among TCR-T cells. Genes with adjusted *p* < 0.01 and Foldchange > 2 were defined as DEGs. Then, signaling pathways involved by these DEGs were estimated by GO analyses relying on the function “compare Cluster” (fun ="enrichGO,’’ pvalue Cut off = 0.1, pAdjust Method = "BH’’, OrgDb = “org.Hs.eg.db,ontBP”) of ChIPseeker R package (v.1.22.1), and visualized by R package “clusterProfiler” ^11^. The cell cycle analysis and regression were performed with the function “CellCycleScoring” and “ScaleData” in Seurat.

**Table S1.** Materials

| **Reagent** | **Supplier** | **Catalog number** |
| --- | --- | --- |
| Celltracker, green CMFDA dye | Invitrogen | C7025 |
| Dulbecco's Modified Eagle Medium (DMEM) | Gibco | 12430054 |
| Dulbecco's phosphate-buffered saline (DPBS) | Gibco | C14190500BT |
| Droplet Generation Oil for EvaGreen | BIO-RAD | 1864005 |
| FEP Tubing,1/16 inch×0.25mm, 10m | DOLOMITE | 3200063 |
| Fetal bovine serum (FBS) | Gibco | 192-1005PJ |
| F68 | Pluronic | 24040-032 |
| Ficoll-PM400 | Cytival | 17030010 |
| Micro tubing (inner dimensions: 0.38mm, outer dimension: 1.09mm) | Scientific Commodities | BB31695-PE/2 |
| Negative photoresist SU-8 | Microchem | SU-8 3050 |
| Penicillin-Streptomycin (P/S) | Gibco | 15140-122 |
| Polydimethylsiloxane (PDMS) | Dow Corning | 02085925 |
| Qubit dsDNA HS assay kit | Invitrogen | Q32851 |
| RPMI1640 medium | Gibco | C22400500BT |
| Silicon (4 inch) | Zhejiang JRH Technology Co., Ltd. |  |
| Single-cell RNA sequencing kit | BGI |  |
| Tris(3-hydroxypropyl)phosphine (THPP) | Sigma-Aldrich | 777854 |
| 1H,1H,2H,2H–perfluoro-1-octanol (PFO) | Sigma-Aldrich | 370533 |
| 2 mL collection tube | Corning | 430659 |
| 1ml syringe | AndeMed |  |
| 10ml syringe | AndeMed |  |
| 30ml syringe | BD | 302833 |

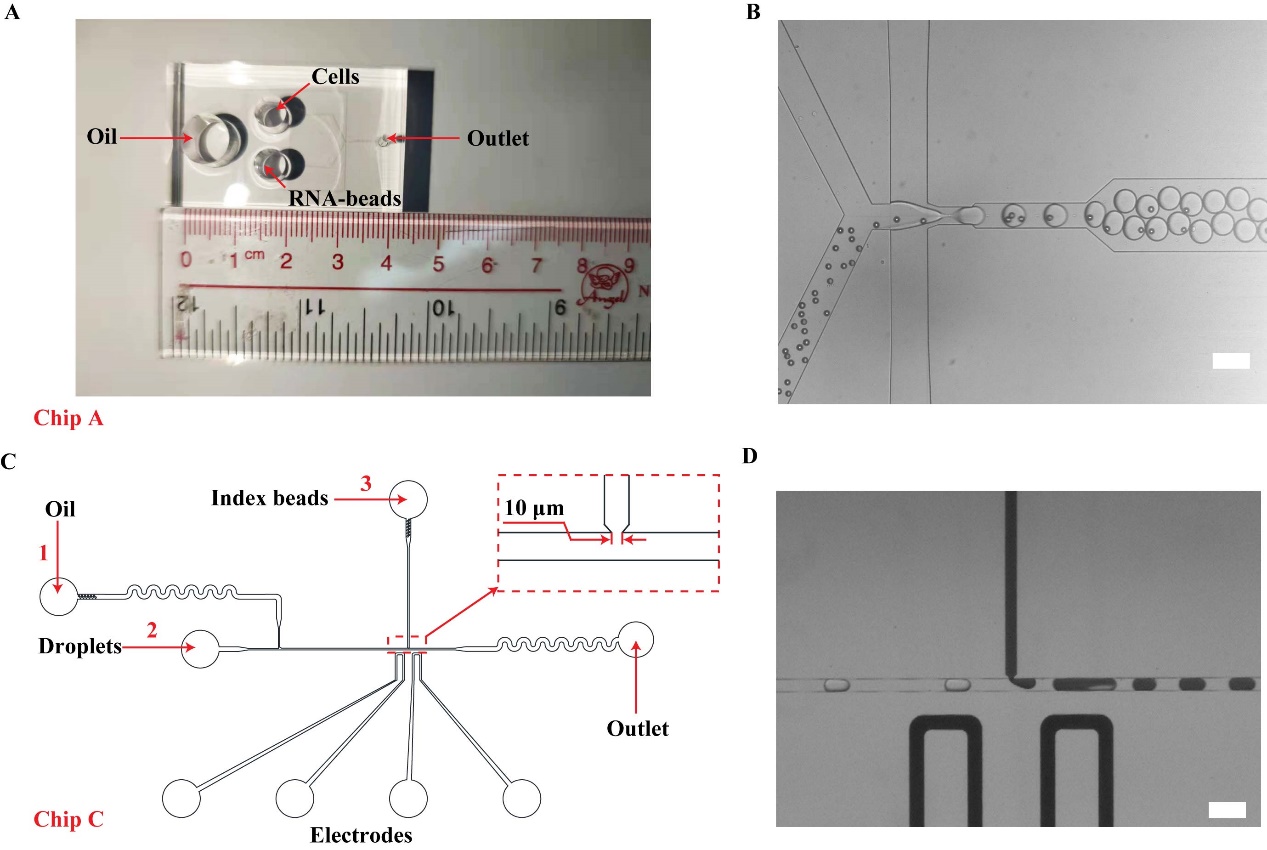

**Fig. S1.** The detailed design of microfluidic chips. A) Design schematic diagram of droplet generation chip (Chip A) and the amplified view of local details; B) Real-time recording of droplet generation by negative pressure. Scale bar: 100 μm; C&D) Design schematic diagram and photograph of droplet injection chip Scale bar: 100 μm.

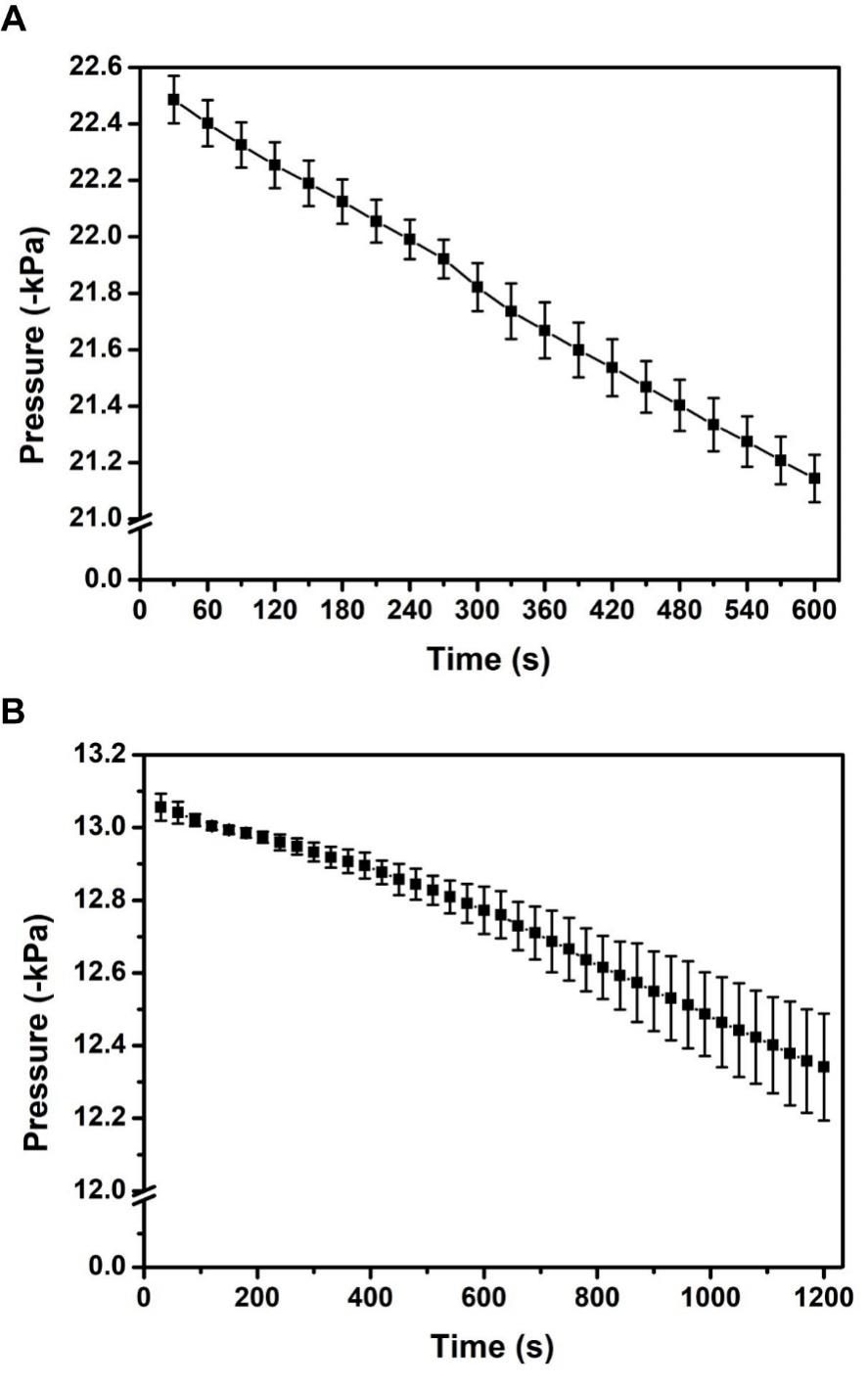

**Fig. S2.** Stability test of negative pressure on microfluidic chips. (A: Chip A; B: Chip C) N=3.

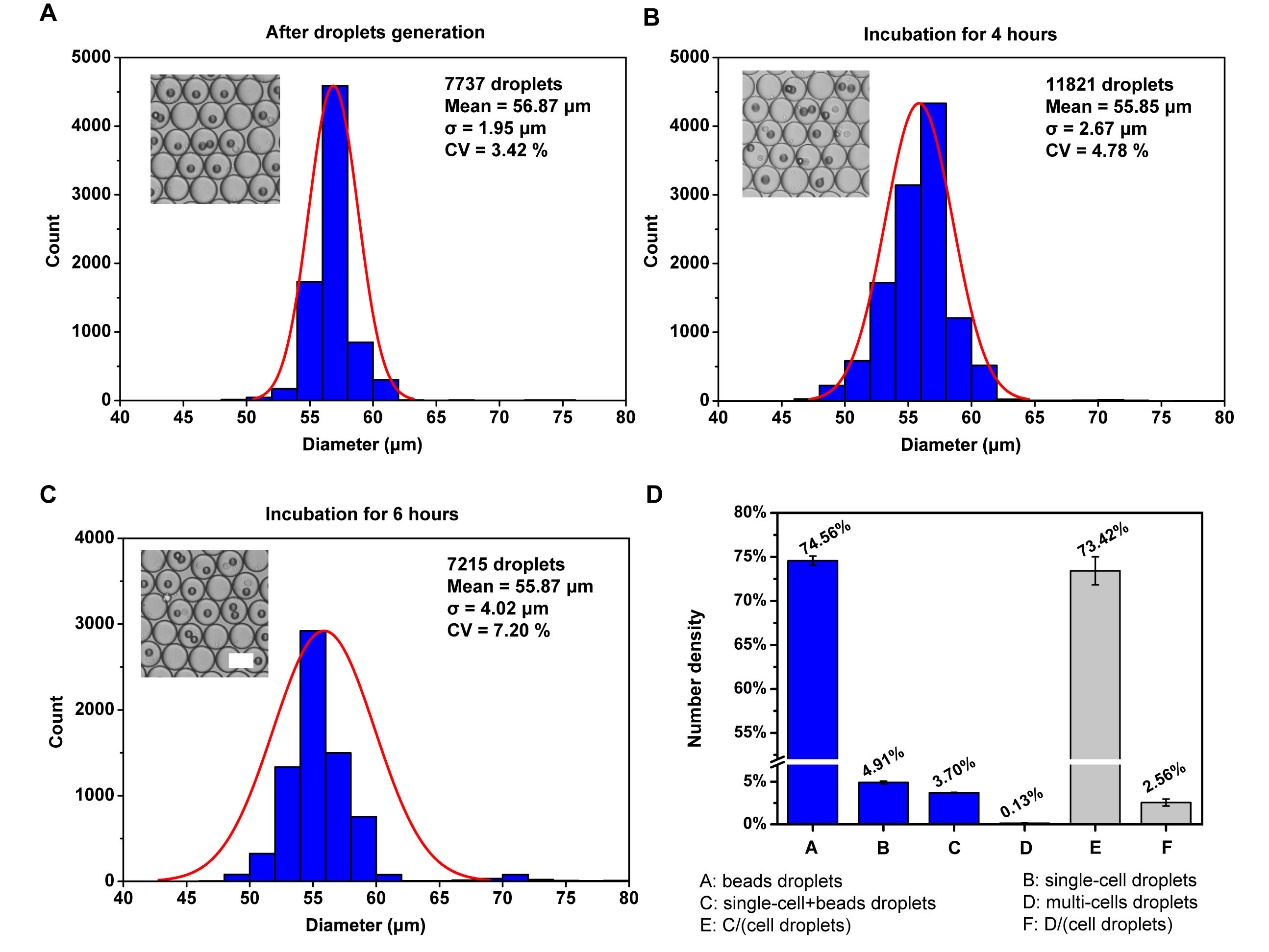

**Fig. S3.** Statistics of the droplet diameter and encapsulation rate of RNA-beads. A) The diameter distribution of 7737 droplets after droplets generation. B) The diameter distribution of droplets after 4 hours incubation. C) The diameter distribution of 7215 droplets after 6 hours incubation. Scale bar: 50 μm. D) The proportional distribution of different type droplets. N = 17113 droplets from three independent experiments.

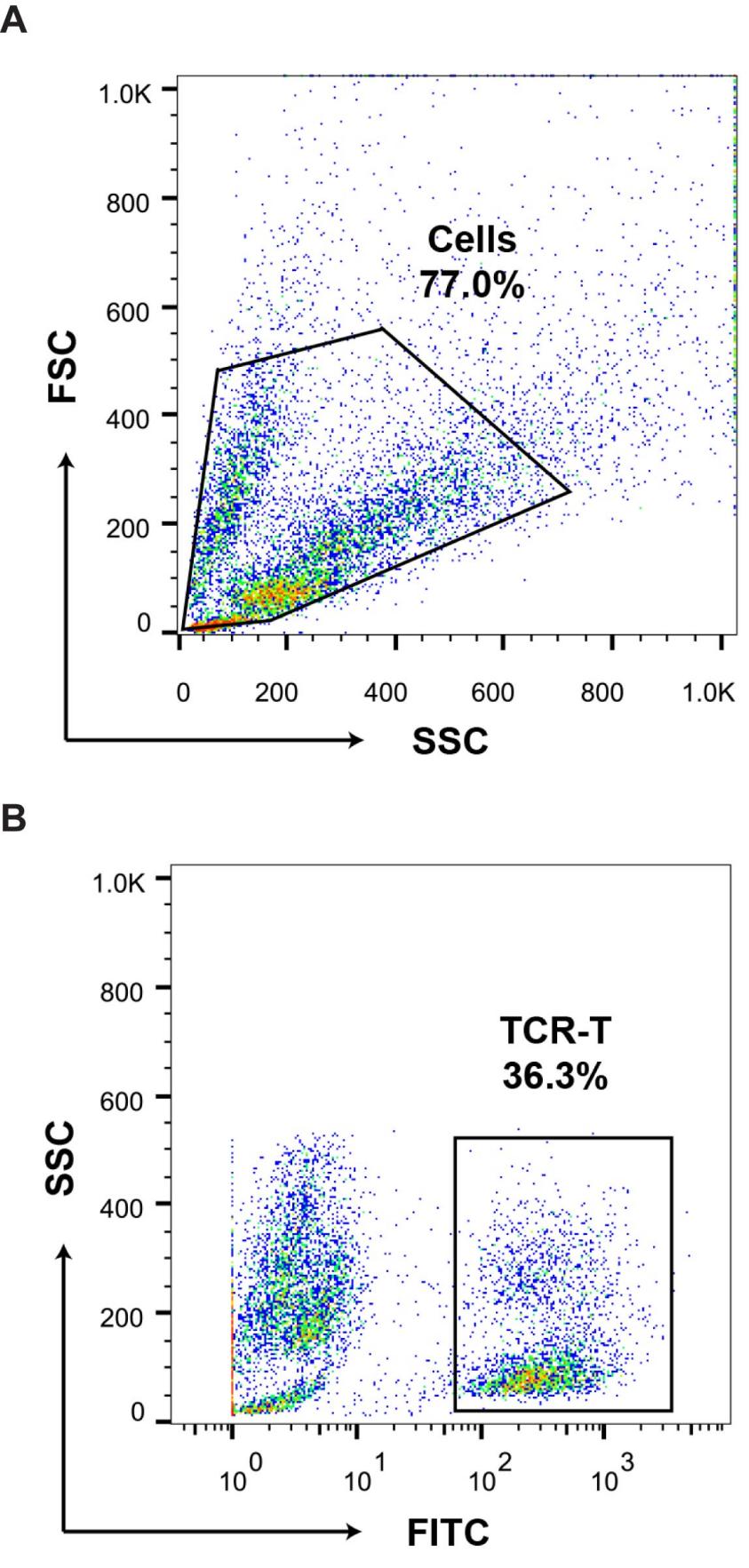

**Fig. S4.** Flow cytometry for the identification of TCR-T cells. A) SSC-FSC plot of cells. B) Gating strategy of TCR-T cells based on the CellTracker CMFDA fluorescence. SSC: side scatter; FSC: forward scatter.

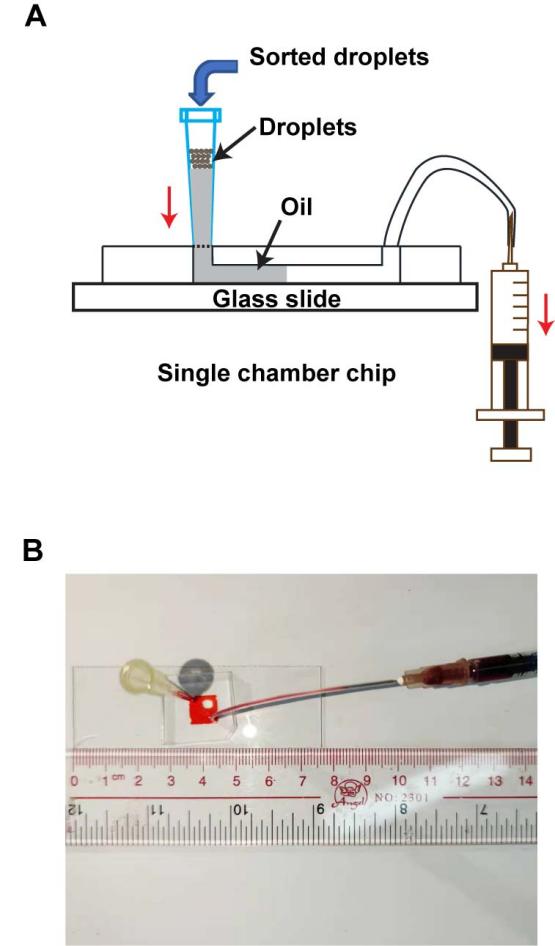

**Fig. S5.** Sorted droplet collector. A) Schematic diagram of the droplet collector. B) A picture of the droplet collector.

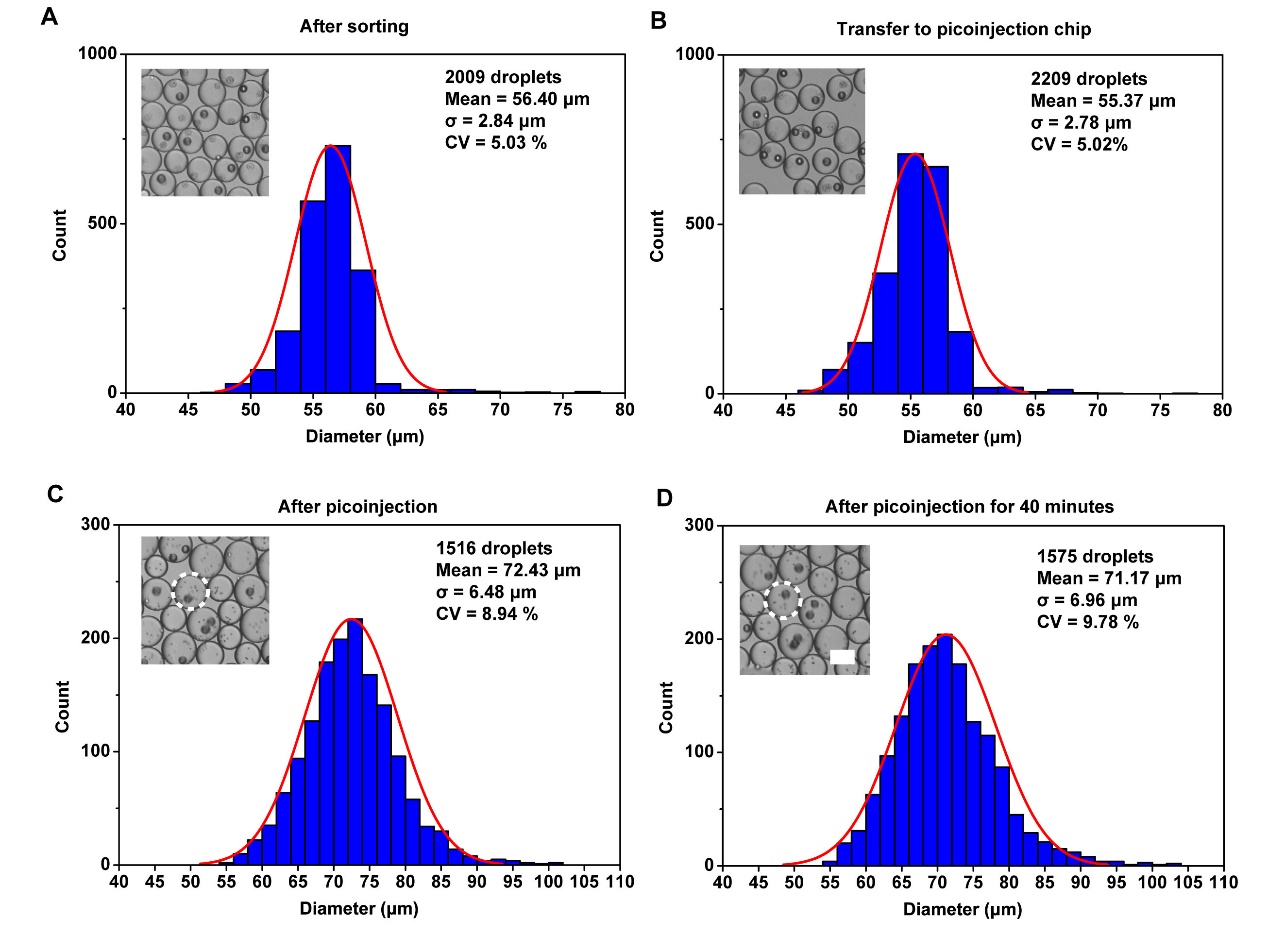

**Fig. S6.** Statistics of the droplet diameter on different conditions. A) The diameter distribution of 2009 droplets after sorting. B) The diameter distribution of 2209 droplets after transferring to picoinjection chip. C) The diameter distribution of droplet after picoinjection. The droplet injected by lysis buffer is circled. D) The diameter distribution of 1575 droplets after picoinjection for 40 minutes. Scale bar: 50 μm.

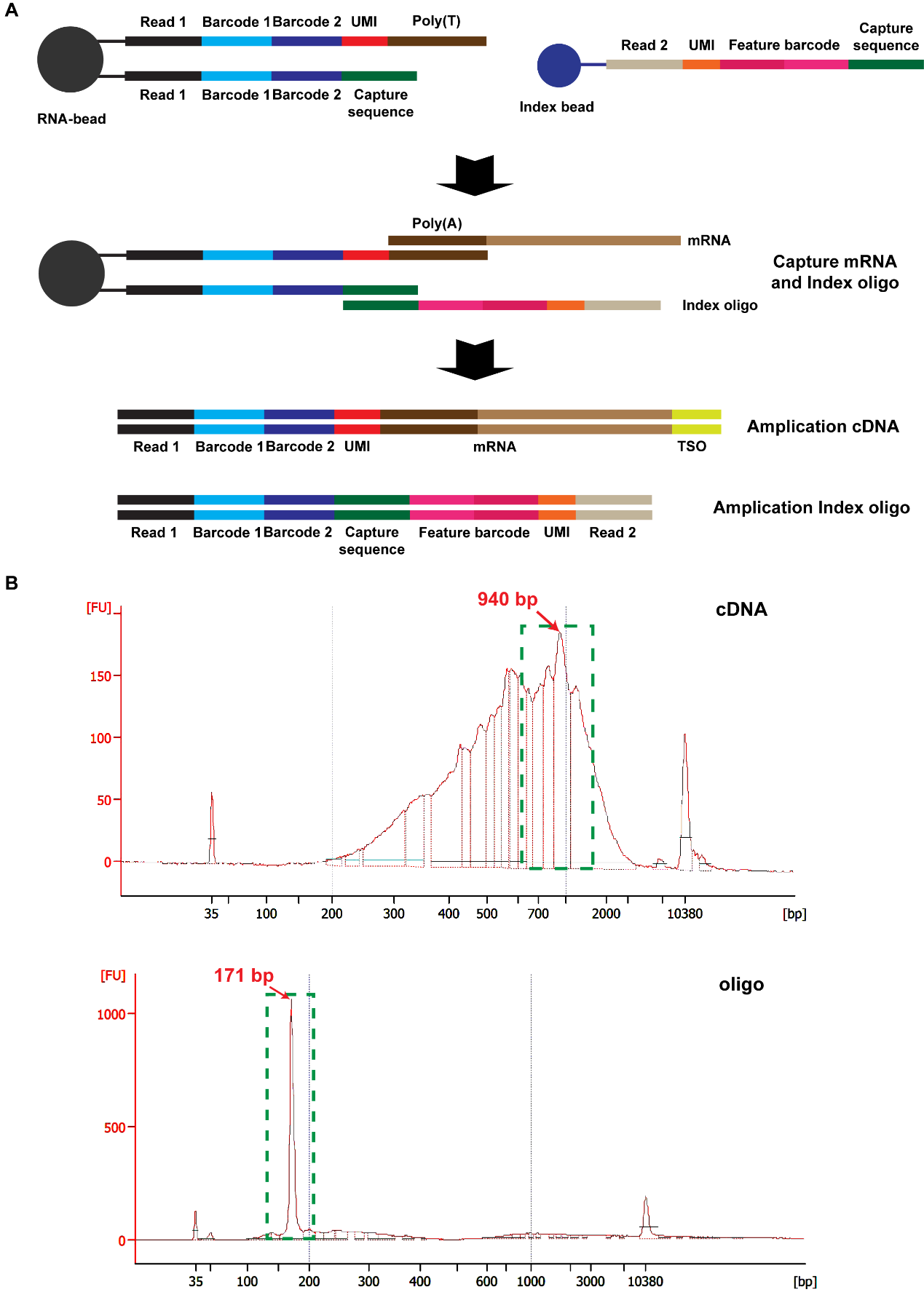

**Fig. S7.** Principle and verification of nucleic acid capture. A) Schematic diagram of library preparation includes capture mRNA & index oligo, and amplify cDNA & index oligo. There was a disulfide bond between index bead and Read2, and the linker could be cleaved by tris(3-hydroxypropyl) phosphine or dithiothreitol. B) Analysis of PCR production of cDNA and oligo. The main peak in the electrophoretogram of cDNA is ~1000 bp, and the electropherogram of oligo has an independent peak around 170bp.

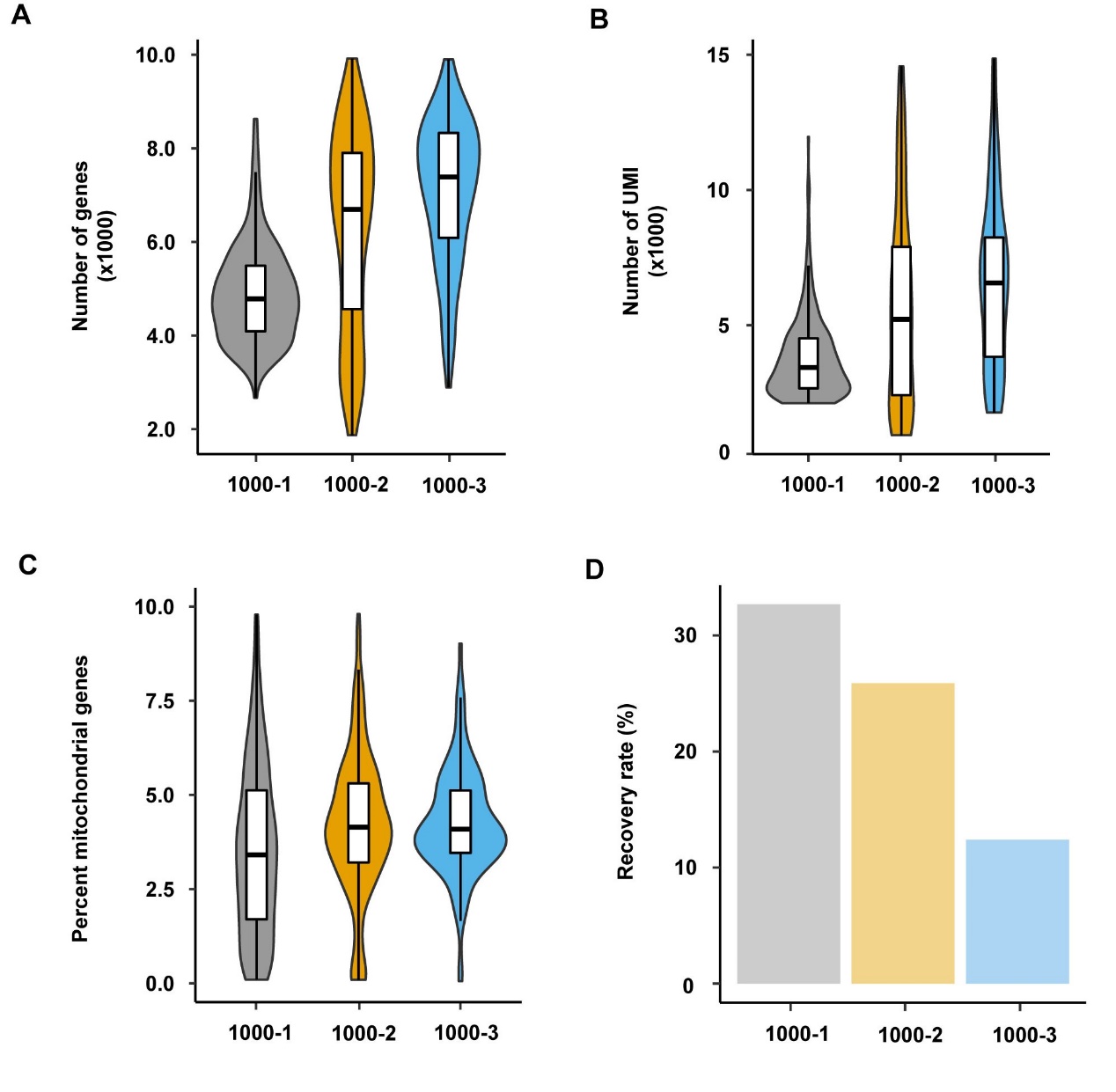

**Fig. S8.** Qualification of 1000-cell samples by RNA sequencing. A) The number of genes per cell for each sample. B) Distribution of UMI number per cell for each sample. C) The percentage of mitochondrial genes per cell. D) The recovery rate of 1000-cell samples.

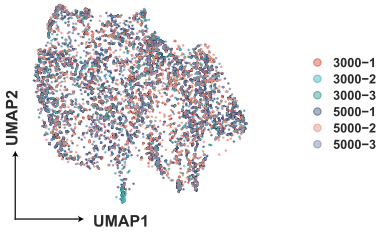

**Fig. S9.** UMAP of cells from different samples. Cells of the six samples were presented on UMAP plot after removing batch effect.

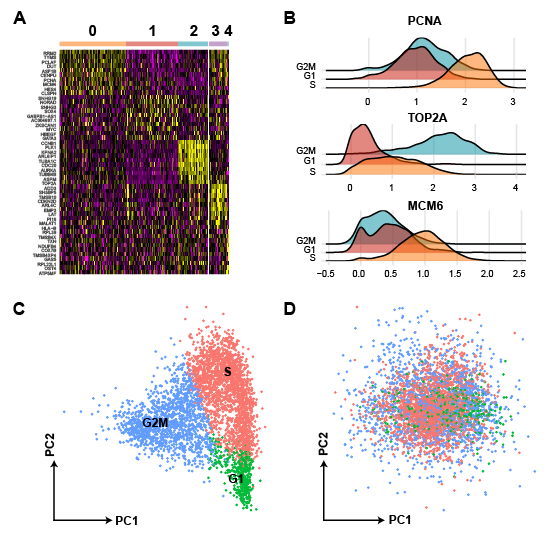

**Fig. S10.** Cell cycle analyses of TCR-T cells. A) Heatmap of top10 feature genes of each cluster. B) Expression of PCNA, TOP2 and MCM6 of cells C) PCA of TCR-T cells at different phases. D) PCA of cells grouped by cell cycle markers with cell cycle genes regression.

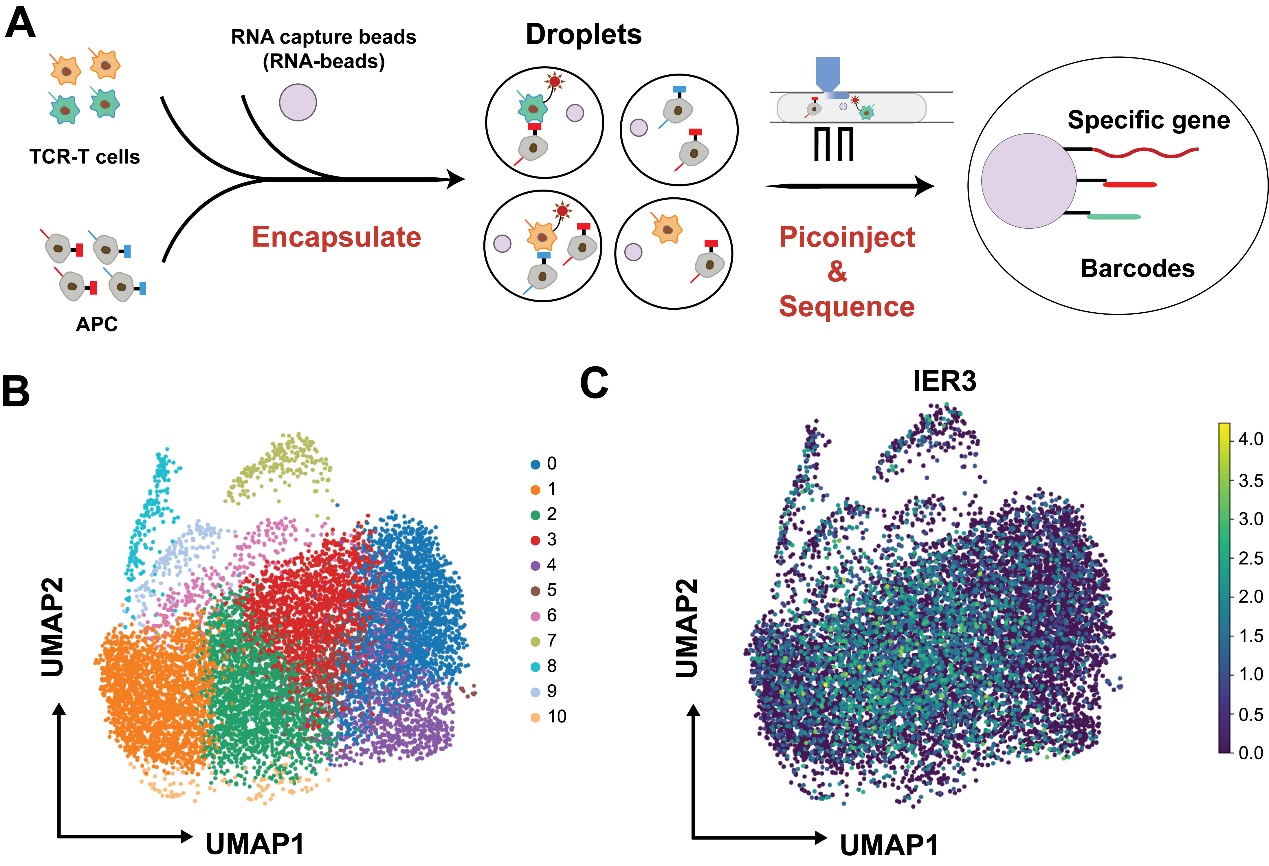

**Fig. S11.** The strategy of next work in antigen-specific TCR recognition. A) A brief schematic diagram of experimental flow. In brief, TCR-T cells and APC cells were labeled with barcodes separately. Next, RNA-beads and cells were wrapped in droplets, and then the reagent injection introduced lysis buffer without destroying the pairing relationship of TCR-APC. After that, the recovery of molecular information, the library generation as well as the data analysis were used to confirm the corresponding barcodes of TCR-T and APC cells. Finally, the effect of TCR screening can be determined by a combination of barcodes and gene expression after the stimulation. B) UMAP of all cells in droplets after picoinjection. C) UMAP of IER3 involved in TCR activation.
